# Adaptation to yeast restriction through the evolution of resource acquisition, macronutrient reserve, and ovarian function

**DOI:** 10.1101/2024.08.24.609539

**Authors:** Purbasha Dasgupta, Anish Koner, Rabi Sankar Pal, Pranav Nandan Pradhan, Kingkini Roychoudhury, Bodhisatta Nandy

## Abstract

Optimization of reproduction under dietary adversity is an important aspect of diet-dependent adaptation. Yet, little is known about the mechanism of such adaptive evolution. Here, we investigated a set of experimentally evolved populations of *Drosophila melanogaster* where early-life fecundity evolved as an adaptation to chronic protein restriction. We tested the role of resource acquisition and macronutrient storage, and changes in ovarian function that could have allowed such reproductive adaptation. We show that higher early-life fecundity was associated with the increased larval feeding rate, aiding in accumulation of higher protein content at the time of eclosion. Further evidence also suggests increase in reproductively allocated lipid content. Evolved females were found to have larger ovaries that also had a higher number of mature, post-vitellogenic oocytes that seem to readily allow the attainment of higher fecundity. Our results further show the evolution of plasticity in ovariole count (depending on mating status) and total egg-chamber count in females. These results are important in understanding the putative role of genetic variance and covariances that aid or limit the evolution of reproductive optimization, especially under nutritional adversity.

## Introduction

Natural selection should lead to the optimization of reproductive investment since resource availability is never unlimited (Kirkwood, 1977; Roff, 1992; Stearns, 1989, 1992, 2000).Allocation in reproductive traits often results in reduced survival and/or reduced future fertility due to constrained resource availability to growth, immune function, and tissue repair (Stearns, 1992). Reproductive optimization is thus pivotal to successful life history strategies. Pianka (1976) conceptualized organisms as “input-output systems” where foraging tactics represented the input of resources, and progeny number represented the output. Thus, it is important to integrate foraging and resource allocation with life history to better understand adaptive reproductive optimization (Pianka, 1976; Boggs 1992; Gatto, Matessi & Slobodkin, 1989).Putatively important role of dietary ecology in reproductive physiology and strategy have not been fully explored beyond the literature on dietary restriction leading to temporary shutting down of reproduction (Mair et al., 2004; Carey et al., 2008; Inness & Metcalfe, 2008; Adler & Bonduriansky, 2014; Moatt et al., 2016; Zajitschek et al., 2016; Zajitschek et al., 2019). Since metabolic energy serves as the currency that links foraging to optimal reproduction, investment is often physiologically regulated (Boggs, 1992; Ellison, 2003; Jervis et al., 2008). Further, the quantity and timing of reproductive allocation have profound implications for lifetime reproductive success (Long et al., 2010; Desprez et al., 2017; Churchill et al., 2019). However, genetic variance and covariances that aid or limit the evolution of reproductive allocation and its timing are poorly understood.Dietary restriction (DR), a well-known intervention, can be a key experimental paradigm to probe resource allocation and reallocation. DR is known to limit reproductive allocation, often resulting in an increased investment in stress resistance and survival (Mair et al., 2005; Partridge et al., 2005; Nakagawa et al., 2012; Fontana & Partridge, 2015). Theoretically, such resource reallocation may enable organisms overcome transient (nutritionally) challenging periods by prioritizing essential functions for immediate survival (Shanley & Kirkwood, 2000; Adler & Bonduriansky, 2014; Zajitschek et al., 2019; Moatt et al., 2020). Interestingly, long-term (multi-generational) nutritional restriction can result in the evolution of strategies aimed at optimizing reproduction (Williams, 1957; Bonduriansky et al., 2008; Zajitschek et al., 2019). Such evolutionary responses can be used to infer on the genetic variance and covariances pertaining to resource acquisition, utilization and storage traits underlying adaptive reproductive optimization (Chippindale et al., 1996; Schwasinger-Schmidt et al., 2012; Ahmad et al., 2018).

Experimental evolution under a variety of dietary conditions has been known to result in adaptations resulting in improved survival and/or reproduction in fruit flies (Kolss et al., 2009; Zajitschek et al., 2016; Cavigliasso et al., 2020). Similar to several other organisms, phenotypic response to short-term nutritional deprivation in *Drosophila* involves a shift from reproduction to survival mode (Rion & Kawecki, 2007; Ballard, Melvin & Simpson, 2008; Lee & Jang, 2014; Shu et al., 2022). For instance, protein restriction caused by yeast limitation has been shown to increase longevity and starvation resistance at the cost of fecundity (Chippindale, 1993; Zimmerman et al., 2003; Mair et al., 2005; Piper et al., 2005), indicating physiological trade-off between egg production and somatic storage, especially lipid reserve (Simmons & Bradley, 1997). Further, in response to a low protein: carbohydrate diet, adult *Drosophila melanogaster* showed an increase in lipid reserve accompanied by greater resistance to starvation stress (Lee & Jang, 2014). Importantly, in laboratory and wild populations, *Drosophila* females largely depend on yeast as the primary protein source, which has a profound impact on reproductive output.Restricted access to yeast, especially live-yeast supplement, appears to have a more targeted effect on decreasing female fecundity, and extending lifespan (Chippindale et al., 1993; Nandy et al., 2012; Magwere et al., 2004). However, such reallocation of resources from reproduction to survival, is unlikely to be adaptive under long-term nutritional restriction. Generations under yeast-deprivation have been shown to lead to the evolution of higher early-life fecundity in experimental populations of *D. melanogaster* (Dasgupta et al., 2022). Interestingly, such reproductive evolution seems to be a readjustment of reproductive schedule, wherein increased early-life fecundity evolved with faster reproductive senescence such that cumulative life-time fecundity and lifespan were unaffected (Dasgupta et al., 2022). Thus, it is amply clear that selection can act on variation affecting early-life fecundity, driving such reproductive adaptation. However, it is not yet clear whether the underlying physiological mechanism of such adaptation involves evolution of resource acquisition (larval or adult) or macronutrient metabolism (for example, storage) or both.

Here, we addressed this unsolved problem by investigating the populations used by Dasgupta et al. (2022). Theoretically, there are two non-mutually exclusive ways of achieving higher early-life reproductive output in females, without an associated survival cost -(a) increased larval resource acquisition (juvenile resource acquisition theory), and (b) increased macronutrient reserve that can cater to the increased metabolic energy demand of higher reproduction (macronutrient content theory). Once the metabolic resources are made available to the female reproductive system, the system can either ramp up the production of eggs and/or increase the number of egg-producing units. Since in holometabolous insects such as fruit flies, early-life fitness heavily depends on larval resource acquisition, we compared early third-instar larval feeding rates across evolved and control regimes to test juvenile resource acquisition theory.

Considering the role of protein and lipid in egg production (Drummond-Barbosa & Spradling, 2001) as well as in survival and stress resistance (Chippindale, Chu & Rose, 1996; Rehman & Varghese, 2021), we compared the whole-body protein content and lipid content in females of evolved vs. control lines to test the macronutrient content theory. Further, we also investigated the putative role of changes in ovarian function by measuring ovarian size (ovariole count), which is known to be a strong determinant of reproductive potential (Wayne et al., 2006; Lobell et al., 2017), and the abundance of mature stages of oocytes in each ovariole. We hypothesized that the increased early-life fecundity in evolved females should be ascribed to either an increase in ovariole count and/or a surge in the production of the number of oocytes per ovariole.

## Materials and methods

### Study populations and selection regime

All the experiments described here were conducted using the four replicate experimental populations (YLB_1-4_) and their corresponding control populations (BL_1-4_). The details of these populations are discussed in Dasgupta et al. 2022. In short, experimental populations (YLB_1-4_) were derived from four replicate baseline populations (BL_1–4_) of *D. melanogaster*, which are outbred (N ∼2800), wild-type populations. BL populations were maintained under a 14-day discrete generation cycle at 25°C on a banana-barley-sugarcane jaggery-yeast diet. Adults are given a supplement of *ad-lib* yeast-paste smeared on food plates. Every generation, eggs are collected during an 18-hour window of the day14 of the generation cycle to start the next generation. This 18-hour window constitutes the (Darwinian) fitness window as in this system, progeny produced only during this window can be passed on to the next generation. Hereafter, “fitness window” refers to this time period of the maintenance regime.

To create YLB populations, ∼2800 eggs were collected from each BL population, and these populations were maintained identically to BL populations, except the yeast supplementation step for adults. Thus, YLB populations are maintained in absence of yeast-supplement at the adult. Prior to assays, all populations were passed through one generation of common-garden rearing conditions to equalize the putative role of parental diet. During this common-garden rearing phase, both BL and YLB populations were provided with *ad-lib* yeast-paste, matching the ecology of BL regime exactly. Wherever needed virgin flies were collected within 6 hours of eclosion under light CO_2_-anaesthesia. A total of six experiments were performed. Experiments 1-3 were conducted within 67-74 generations; experiment 4 on the 79^th^ generation; experiment 5 within the 94-95 generations and finally, experiment 6 on the 120^th^ generation of the experimental evolution.

#### Experiment 1: Whole-body protein content in females

The general plan was to measure protein content at two time points – on eclosion, and during the fitness window (i.e., day14 of the generation cycle) in reproducing condition. In this assay, one set of flies was used to measure whole body protein content using the Bradford method (see Supplementary information S1). The other set was used to measure whole body mass, which is used as a scaling factor at the time of statistical analysis.

For protein quantification on eclosion, females were collected as virgins in two sets and immediately frozen at -20°C. In one set, groups of three females were collected in an empty vial, which were processed for protein content measurement within 24 hours. A second set of females were collected in groups of five, which were processed for body weight measurement within 24 hours. Twelve such vials were measured per population. For body weight measurement, flies were dried in a hot air oven for 24 hours at 30°C (following Colinet & Renault, 2014) and weighed using Shimadzu AUW220D to the nearest 0.01 mg in the pre-formed groups. Each group of five females was weighed thrice and the average of the three readings was noted. A population mean was calculated using values from all replicate vials. These population means were used to normalize the protein content value obtained from the Bradford assay (see Supplementary information S1). For protein quantification in the fitness window, adult flies were collected as virgins and held in single-sex food vials in groups of ten. 10 such vials were set up for each sex per population. On day12 post-egg collection, 2-3 days old virgin males and females were transferred into fresh food vials and were allowed to interact for 48 hours during which they completed mating. Afterwards, the vials containing flies were frozen at -20°C. Frozen females were sorted in groups of three in empty glass vials. These females were later used for protein quantification using the Bradford assay (see Supplementary information S1). Here, each group of three females served as the unit of replication. Additionally, 12 vials with five females in each were kept for dry weight measurement. The latter was done within 24 hours of freezing, following the same protocol described above.

#### Experiment 2: Whole-body dry weight and lipid content in females

The general plan was to measure whole body dry weight and lipid content in freshly eclosed females and in reproducing females at the fitness window. For the first time point, i.e., on eclosion, virgin females were collected in groups of 10 and held in a glass vial. 12 such vials were collected for a population and were immediately frozen at -20°C. Within 24 hours of freezing, all the flies were transferred to a hot air oven and allowed to dry for 48 hours at 60°C before being processed and weighed.

For the second time point, adult flies were collected as virgins and allowed to pass through the following steps before being subjected to the body weight and lipid content measurement.

Freshly eclosed flies were sexed and held in single-sex vials in groups of 10 until day12. For a population, 12 such vials were set up per sex. On day12 post egg collection, males and females were transferred into fresh food vials, and were allowed to interact for 48 hours, following which males were discarded and females were frozen at -20°C. The frozen dead flies were sorted into groups of 10 females and 12 such vials were set. Within 24 hours of freezing, all the flies were transferred to a hot air oven and allowed to dry for 48 hours at 60°C before being processed and weighed.

Dried females were weighed in groups of 10 using Shimadzu AUW220D to the nearest 0.01 mg. Each set of 10 females is weighed thrice and the average across these three measurements were used as Dry Mass (DM) value. After that, each set of 10 individuals was transferred to 1.5 ml centrifuge tubes with 1.5ml Diethyl ether (Marron et al. 2003) and incubated in an incubator shaker at 200 RPM for 24 hours at room temperature (∼25-30°C). After 24 hours, the centrifuge tubes were kept in a hot air oven, with the lids open, at 60°C for another 24 hours to allow the Diethyl ether to evaporate. Following this, each set of 10 individuals was weighed thrice again and the average across these three measurements was obtained. By subtracting this value from the DM value previously obtained, an Absolute Lipid Mass (ALM) value was obtained for each set of ten females. Thereafter, relative lipid content per unit dry mass was calculated by using the formula mentioned below:

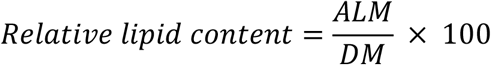

This was done for all twelve samples from a population. This lipid extraction method using Diethyl ether integrates all lipid classes and does not distinguish between lipid sequestered in eggs and that found in somatic tissues (Ballard et al. 2008). From one round of lipid estimation experiments, three sets of data were obtained -dry mass, absolute lipid mass and relative lipid content (i.e., lipid content per unit dry mass). For the final analysis, after the exclusion of outliers, based on 1.5 IQR assessment, values from 10-12 samples were considered for each population.

#### Experiment 3: Starvation resistance

Starvation resistance was assayed by quantifying longevity of the females of both regimes under starved condition. Like other assays mentioned above, the populations were passed through one generation of common rearing following which eggs were collected under standard density. On eclosion, flies were collected and held in single-sex vials in groups of 12. On day 3 post-eclosion, the ‘interaction vials’ were set up by combining 12 males and 12 females in a food vial. 12 such vials were set-up for each population. The vials were left undisturbed for 2 days, following which the females from each vial were separated under light CO_2_ anaesthesia and transferred to ‘starvation vials’. These starvation vials were standard 8-dram vials provisioned with non-nutritive agar gel (1% Agar-agar solution + 1% p-hydroxy benzoic acid solution). The Agar-gel provided moisture for the flies and ensured that flies are only starved but not desiccated. Mortality was recorded in each starvation vial at 6-hours intervals. The mean longevity of 12 individuals per vial under starved condition provided the measure of starvation resistance and was used as the unit of analysis. We removed a few vials from the analysis due to a variety of handling errors, including escaped flies and broken vials, during the assay. The final number of vials used for the analysis ranged from 11 to 12 vials for a population.

#### Experiment 4: Ovariole count

After one generation of common garden rearing, eggs were collected from commonly reared cages and cultured following the standard conditions. On eclosion, adult flies were collected as virgins every 6 hours. Virgin males and females were collected in groups of 10 and held in single-sex vials. These vials were left undisturbed for two days. On day 3 post eclosion, flies were transferred into fresh food vials, and these vials were randomly divided into two (mating status) sets – (a) Mated: 10 males and 10 females placed in a vial and (b) Virgin:10 virgin females placed in a vial. These vials were left undisturbed for two days.

After two days of conditioning, the flies were frozen at -20°C. The number of ovarioles were counted in both the ovaries. This was done on 15 randomly picked females per population for both virgin and mated. Ovaries were dissected out on a drop of PBS and stained with a saturated solution of potassium dichromate for 10-20 minutes, which facilitated the easy identification and counting of ovarioles (Coyne et al., 1991). The ovariole number of each female was defined as the mean of the number of ovarioles in the right and left ovaries. The mean ovariole count of each female was used as the unit of analysis.

Before dissection, thorax length of the females was measured as a proxy for body size to look for possible correlations between body size and ovariole count. Lateral view images of the thorax were captured using *ZEISS AxiocamERc 5s* mounted on a stereozoom microscope. Thorax length was measured as the distance between the highest point of the anterior tip of the thorax and the posterior-most point of the scutellum. The image analysis was done on *Zeiss ZEN 2 Blue edition* software. Each individual was measured thrice, and the mean was used for the analysis.

#### Experiment 5: Larval feeding rate

After common garden rearing, two small food plates were introduced in each population cage and the females were allowed 1 hour for oviposition to ensure developmental synchrony of the experimental larvae. Following the protocol by Joshi and Mueller (1996), about a hundred eggs were then randomly picked from the two oviposition plates and gently transferred to two Petri dishes containing a thin layer of solidified agar gel (1% agar-agar solution) using a fine brush. 24 hours later, 25 newly hatched larvae were gently transferred to a Petri dish containing a thin layer of 1% non-nutritive agar overlaid with 1.5 ml of 37.5% yeast suspension. Four such Petri dishes were set up for each population. They were left undisturbed till the larvae reached an early third-instar stage. 25 larvae from each population were used to quantify the feeding rate by placing them individually on Petri dishes containing a thin layer of agar overlaid with 10% yeast suspension. Each larva was allowed to acclimatize for 15 seconds following which the feeding rate was quantified by counting the number of cephalo-pharyngeal sclerite retractions per minute under a stereo zoom microscope (*ZEISS Stemi 508 Stereo Microscope*). Blind-fold observation was adopted to avoid observer bias. Feeding rate of each larva was taken as the unit of analysis. Here and in all subsequent analyses, a few outliers were removed from the final analysis using ‘±1.5×Interquartile range’ rule. Hence, the final number of larvae used for the analysis ranged between 24-25 for each population.

#### Experiment 6: Stage-composition of the ovarioles

In *D. melanogaster*, each ovariole comprises a string of progressively developing egg chambers or follicles. The anterior end of an ovariole is the germarium containing somatic and germline stem cells. Egg chambers bud off from the germaria and mature through 14 different stages as they move towards the posterior end near the oviduct. Stage 1-7 are in pre-vitellogenic, stage 8-10 are in vitellogenic and stage 11-14 are in post-vitellogenic stages of development (Drummond-Barbosa et al. 2001). Protocols for generation of flies and dissection were identical to those followed in Experiment 4. During the assay, the ovarioles were carefully teased apart using stainless steel needles. Stage-specific quantification of the egg chambers was performed on either the left or right ovary, randomly selected from 15 randomly chosen females per population for both virgin and mated groups. The ovaries were dissected in 1X PBS and fixed in 4% paraformaldehyde for 10-30 minutes. Following fixation, the ovaries were washed four times in 1X PBS, with each wash lasting 15 minutes. To increase tissue permeability, 0.1-0.2% Triton-X was added for 10 minutes. The tissues were then washed in PBS for 15 minutes. Subsequently, the ovaries were stained with 1 µg/ml DAPI for 5-10 minutes and washed again in PBS for 5-10 minutes. Next, the ovarioles were carefully teased apart using a fine tungsten needle and mounted with Vectashield antifade mounting media. The total number of egg chambers at each stage was counted for every female using a fluorescence microscope (*Zeiss Axio Scope A1*). The proportion of pre-vitellogenic, vitellogenic, and post-vitellogenic egg chambers was calculated for each female and used as the unit of analysis. The different developmental stages of the egg chambers were identified based on the characteristics described in Jia et al. (2016).

### Data analysis

All analyses were done on R version 4.0.3 (R Core Team 2020). Linear and generalized linear mixed-effects models (LMM and GLMM respectively) were performed using the package lme4 (Bates et al. 2014). Further, to obtain p-values based on the Satterthwaite approximation for denominator degrees of freedom, we used the R package lmerTest (Kuznetsova et al. 2017). Selection regime was modeled as a fixed factor. Block and its interaction with other fixed-effects were fitted as random effects.

Larval feeding rate was analysed as a GLMM with a Poisson distribution. An OLRE (Observation-level random effect) was included in the model to correct for overdispersion. Female whole-body protein content, absolute lipid mass, relative lipid content and longevity under starvation were analysed using LMM. The macronutrient contents (protein and lipid) were quantified at two different adult life-stages/time points -on eclosion and in fitness window.

Initially, time point was fitted as a fixed effect, however, a significant time point × block interaction was observed. Hence, separate analyses were performed to assess the effect of selection regime on the macronutrient content at each of these time points. On the other hand, female dry body mass was analysed using LMM with selection regime, time point and their interaction as fixed factors.

Ovariole count was analysed using an LMM. Selection regime, mating status and their interaction were fitted as fixed effects, whereas thorax length was included as a covariate. The effect of selection on thorax length was analysed using a GLMM with a gamma distribution and a log link function. To analyse the proportion of pre-vitellogenic, vitellogenic, and post-vitellogenic egg chambers per ovary per female, separate models were used. The proportion of pre-vitellogenic egg chambers was analyzed using a LMM, whereas the proportions of vitellogenic and post-vitellogenic egg chambers were analyzed using GLMMs with a Poisson distribution and OLRE to account for overdispersion. In these models, the number of egg chambers from each stage was the dependent variable, scaled by the total number of egg chambers across all stages, included as an offset.

Wherever applicable, post hoc analyses were done using Tukey’s HSD with emmeans package (Lenth et al. 2018). The final models used for the analyses are described in Table S5. Descriptive statistics of all the traits quantified here are mentioned in Table S4.

## Results

### Experiment 1

Analysis of the whole-body protein content results suggested a significant effect of the selection regime in freshly eclosed females. Compared to BL females, YLB females eclosed with about 15% higher protein content per unit dry mass (Figure 1a, Table 1, Table S4). At fitness window, the effect of the selection regime on protein content was not significant (Figure 1b, Table 1).

**Table 1.** Results of linear mixed effect model (LMM) analyses of various traits. All tests were performed considering α= 0.05 and significant p-values are mentioned in bold.

| Traits | Stage | Effects | SS | MS | Den. DF | F | p |
| --- | --- | --- | --- | --- | --- | --- | --- |
| **Protein content** | On eclosion | Selection regime | 16332 | 16332 | 85.01 | 20.68 | **<0.01** |
|  | In fitness window | Selection regime | 737.1 | 737.1 | 2.94 | 0.95 | 0.4 |
| **Dry body mass** |  | Selection regime (SR) | 4808.3 | 4808.3 | 5.78 | 25.55 | **<0.01** |
|  |  | Time point | 20764 | 20764 | 2.99 | 110.33 | **<0.01** |
|  |  | SR × Time point | 1383.4 | 1383.4 | 5.78 | 7.35 | **0.03** |
| **Absolute lipid mass** | On eclosion | Selection regime | 571.21 | 571.21 | 2.99 | 4.93 | 0.11 |
|  | In fitness window | Selection regime | 2332.5 | 2332.5 | 86 | 17.43 | **<0.01** |
| **Relative lipid content** | On eclosion | Selection regime | 59.9 | 59.9 | 2.94 | 8.33 | 0.06 |
|  | In fitness window | Selection regime | 41.84 | 41.84 | 2.96 | 6.32 | 0.09 |
| **Starvation resistance** |  | Selection regime | 117.21 | 117.21 | 3.07 | 3.02 | 0.18 |
| **Ovariole count** |  | Selection regime (SR) | 2.15 | 2.15 | 222.12 | 0.73 | 0.39 |
|  |  | Mating status (MSt) | 25.57 | 25.57 | 222.17 | 8.62 | **<0.01** |
|  |  | Thorax length | 8.22 | 8.22 | 189.93 | 2.78 | 0.09 |
|  |  | SR × MSt | 29.47 | 29.47 | 221.89 | 9.94 | **<0.01** |
| **Pre-vitellogenic egg chambers** |  | Selection regime (SR) | 965.91 | 965.91 | 5.98 | 4.59 | 0.07 |
|  |  | Mating status (MSt) | 320.10 | 320.10 | 3.01 | 1.52 | 0.30 |
|  |  | SR × MSt | 190.08 | 190.08 | 5.98 | 0.90 | 0.38 |

**Figure 1.**
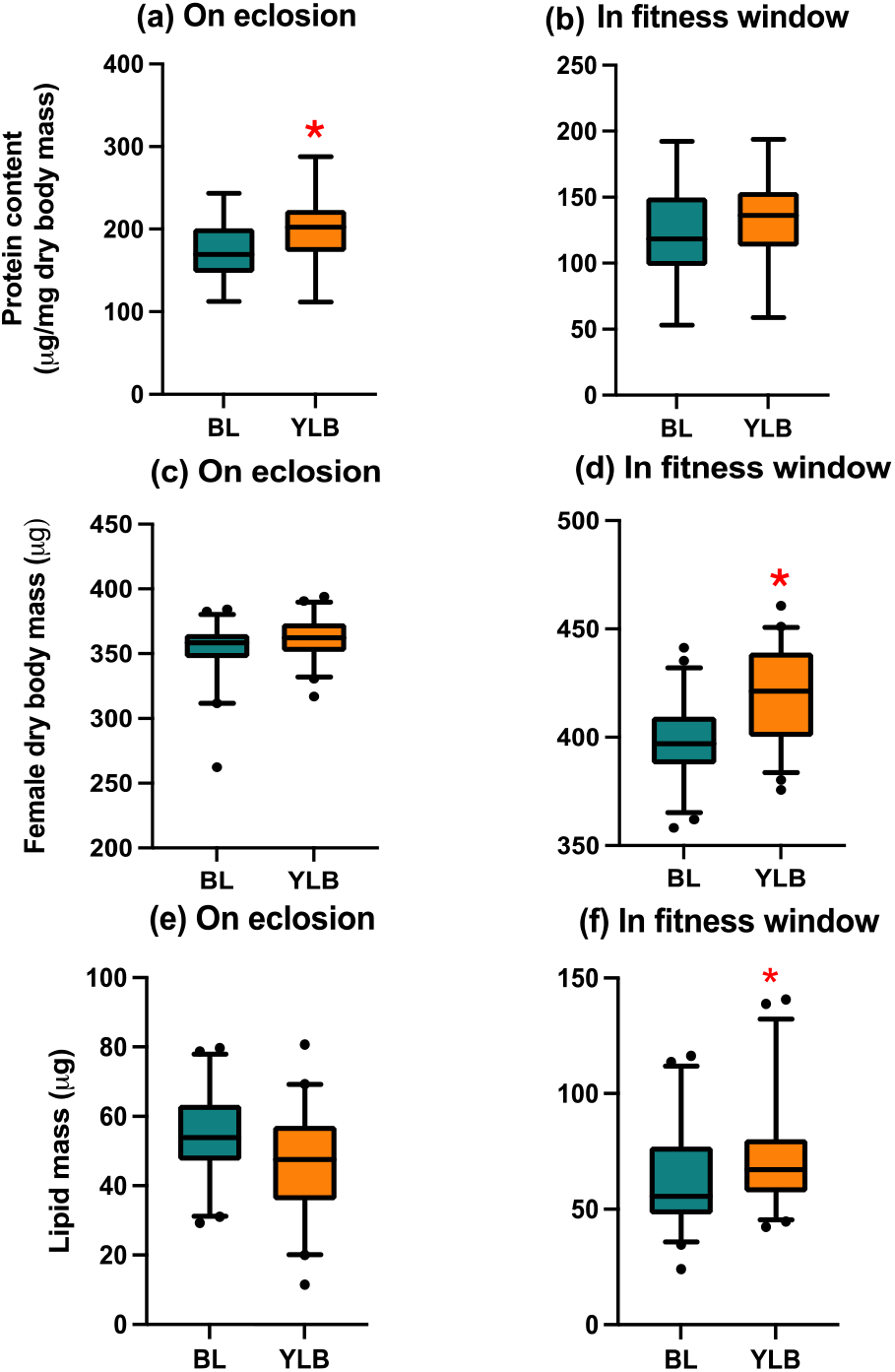
Female whole body protein content (a & b), dry mass (c & d), and lipid content (e & f). Whole body protein content in females from Experiment 1, quantified at two different adult life-stages – on eclosion and in fitness window. Dry body mass from Experiment 2, quantified at two different adult life-stages/time-points – on eclosion and in fitness window. Absolute lipid mass in females from Experiment 3. Whole body absolute lipid mass of females, from Experiment 2, was quantified at two different adult life-stages – on eclosion and in fitness window. The cyan box represents data from BL (control regime) females, while the orange box represents that from YLB (experimental regime). The box thickness represents the Inter-Quartile Range. The black horizontal line in the middle is the median. Wherever the BL vs. YLB difference is statistically significant, it is indicated with an asterisk (*).

### Experiment 2

Analysis of the whole-body dry weight suggested a significant effect of selection regime, time point and the interaction between them (Table 1). On an average, dry mass of the YLB females was significantly higher compared to that of the BL females. Pairwise comparisons suggested that, on eclosion the dry weight difference was not significant, however, at the fitness window, dry weight of the YLB females were found to be significantly higher compared to that of the BL females (Table S1). While females of both the regimes showed a significant increase in dry mass from eclosion to their fitness window, the degree to which it increased appeared to be different. YLB females showed a 16% increase in dry weight whereas, a 12% increase was observed for the BL females between these two time points (Figure 1c,d; Table 1, Table S4).

Analysis of the lipid content measurement indicated that the effect of the selection regime was not significant at the first time point, i.e., on-eclosion (Table 1, Figure 1e). This was true for both measures of lipid content, i.e., absolute lipid content as well as relative lipid content, at this time point. At the second point, i.e., during the fitness window, the effect of the selection regime on absolute lipid content was significant but the same was not true for relative lipid content (Table 1). YLB females were found to have ∼16% higher mean absolute lipid mass compared to the BL females at the second time point (Figure 1f, Table S4). However, this difference turns out to be non-significant after factoring in the dry mass difference. This implied that the observed difference in dry mass between the two types of females at the second time point was primarily due to the difference in lipid content.

### Experiment 3

We did not find a significant effect of the selection regime on females’ mean longevity under starved conditions (Table 1, Figure 2b). This indicated that there was no difference in starvation resistance between the YLB and BL females.

**Figure 2.**
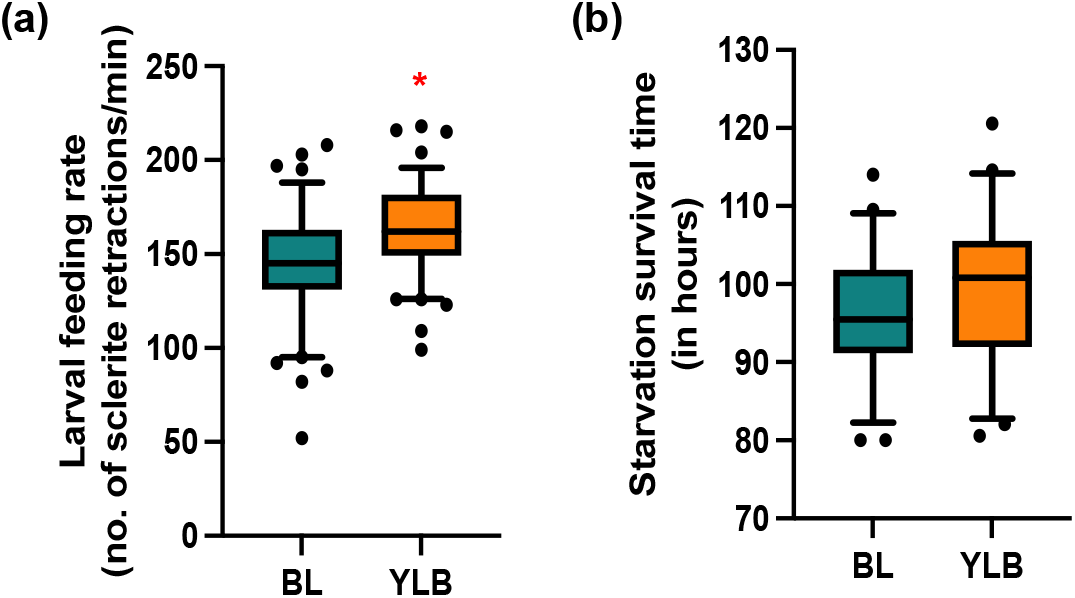
Important results from Experiment 3 and 5. (a) Larval feeding rate (Experiment 5) was quantified as the number of sclerite retractions per min in an early third-instar larva. (b): Starvation resistance (Experiment 3) was quantified as the females’ mean longevity under starvation. The cyan box represents data from BL (control regime), while the orange box represents that from YLB (experimental regime). The box thickness represents the Inter-Quartile Range. Selection regime had a significant effect on larval feeding rate. Wherever the BL vs. YLB difference is statistically significant, it is indicated with an asterisk (*).

### Experiment 4

Analysis of the ovariole count results indicated that the fixed effect of selection was not significant (Table 1). However, the effects of mating status, and selection regime × mating status interaction were significant (Table 1). Tukey’s adjusted P-value obtained by pairwise comparisons suggested that while there was no significant difference in ovariole numbers between BL and YLB females under virgin condition. However, under mated condition, a significantly higher (∼5%) number of ovarioles was observed in YLB females (Table S2, Figure 3). The effect of thorax length, as a covariate, was not significant (Table 1). We also tested the effect of selection regime on thorax length and found the effect to be non-significant, indicating that body size did not respond to experimental evolution (Table 2).

**Table 2.**
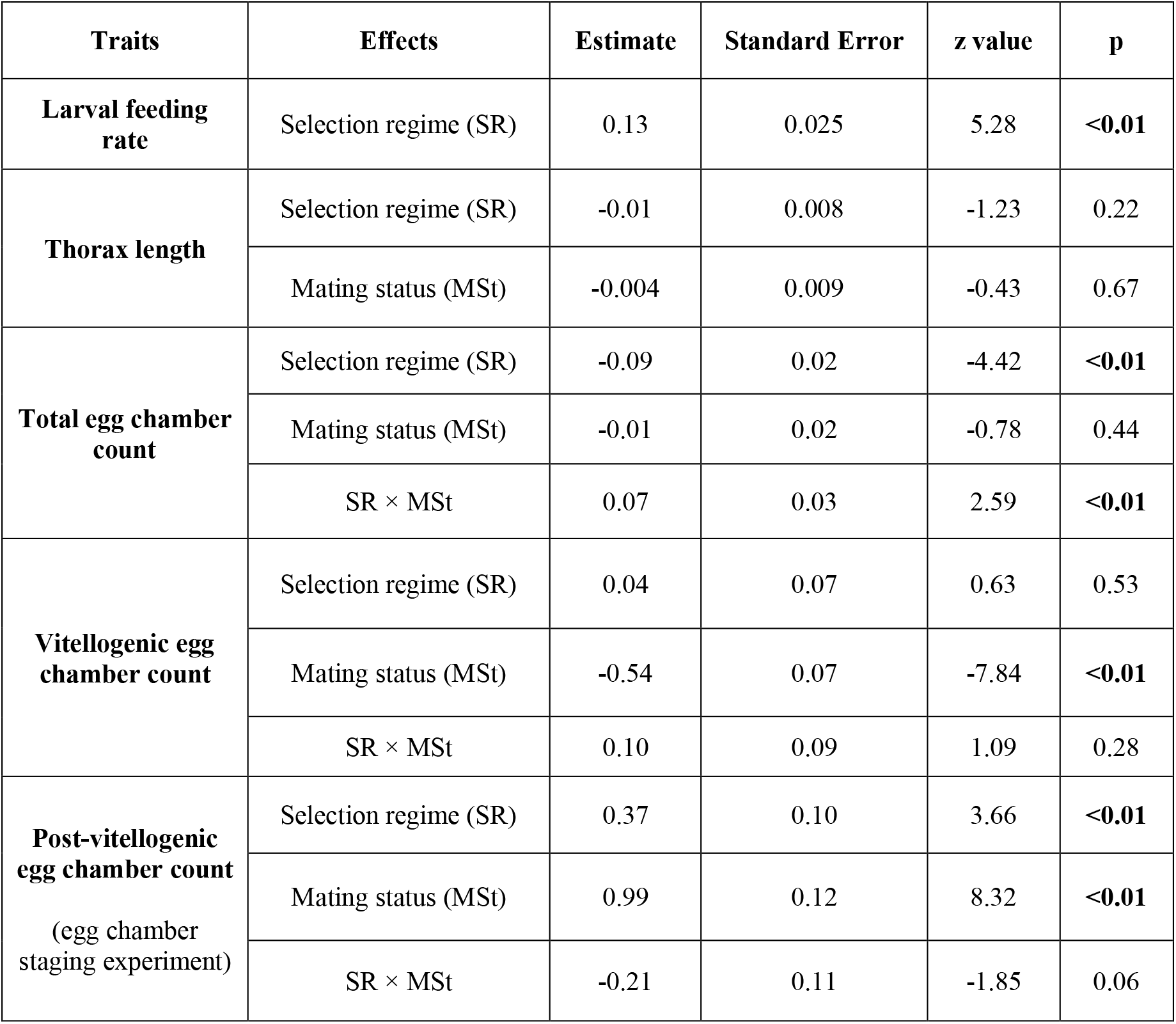
Results of generalized linear mixed effect model (GLMM) analyses of various traits. All tests were performed considering α= 0.05 and significant p-values are mentioned in bold.

| Traits | Effects | Estimate | Standard Error | z value | p |
| --- | --- | --- | --- | --- | --- |
| **Larval feeding rate** | Selection regime (SR) | 0.13 | 0.025 | 5.28 | **<0.01** |
| **Thorax length** | Selection regime (SR) | -0.01 | 0.008 | -1.23 | 0.22 |
|  | Mating status (MSt) | -0.004 | 0.009 | -0.43 | 0.67 |
| **Total egg chamber count** | Selection regime (SR) | -0.09 | 0.02 | -4.42 | **<0.01** |
|  | Mating status (MSt) | -0.01 | 0.02 | -0.78 | 0.44 |
| | SR × MSt | 0.07 | 0.03 | 2.59 | **<0.01** |
| **Vitellogenic egg chamber count** | Selection regime (SR) | 0.04 | 0.07 | 0.63 | 0.53 |
|  | Mating status (MSt) | -0.54 | 0.07 | -7.84 | **<0.01** |
| | SR × MSt | 0.10 | 0.09 | 1.09 | 0.28 |
| **Post-vitellogenic egg chamber count**<br>(egg chamber staging experiment) | Selection regime (SR) | 0.37 | 0.10 | 3.66 | **<0.01** |
|  | Mating status (MSt) | 0.99 | 0.12 | 8.32 | **<0.01** |
| | SR × MSt | -0.21 | 0.11 | -1.85 | 0.06 |

**Figure 3.**
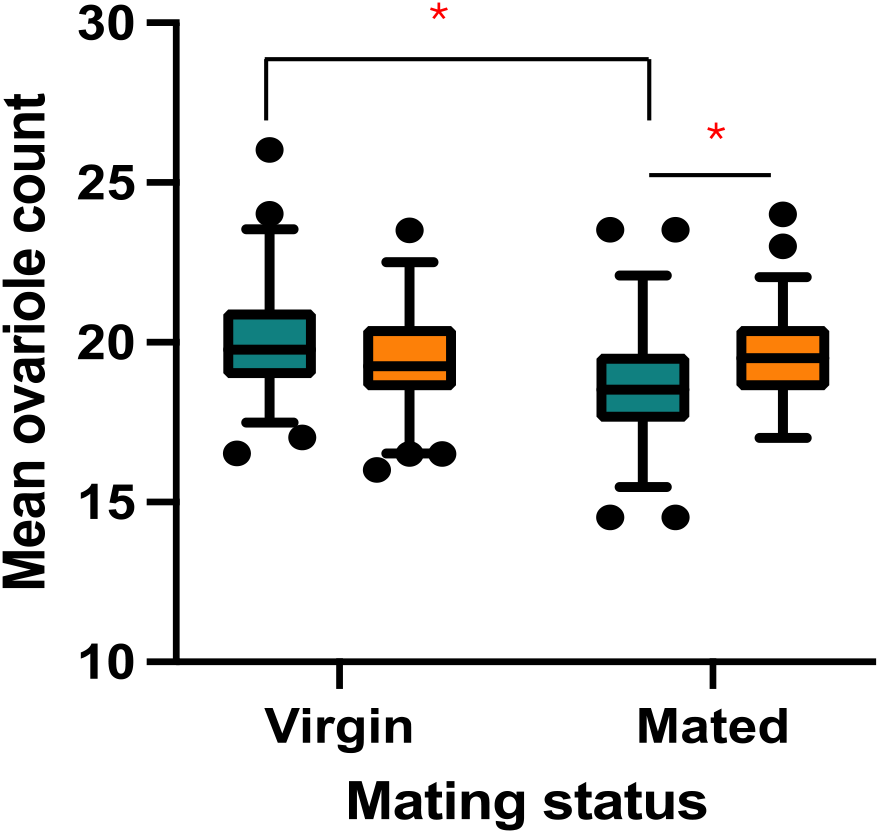
Mean ovariole count from Experiment 4. Ovariole number was quantified for virgin and mated females. The cyan box represents data from BL (control regime) females, while the orange box represents that from YLB (experimental regime). The box thickness represents the Inter-Quartile Range. The black horizontal line in the middle is the median and the notch denotes 95% CI around the median. Statistically significant differences, as indicated by multiple comparisons, are indicated with an asterisk (*).

### Experiment 5

The analysis of the feeding rate of the early third-instar larvae suggested a significant effect of the selection regime (Table 2). The third instar larvae of YLBs were found to have a significantly higher (∼13%) feeding rate compared to those of the BLs (Figure 2a).

### Experiment 6

The analysis of the total number of oocytes per unit ovariole, regardless of developmental stage, revealed significant effects of selection regime and selection regime × mating status interaction (Table 1). BL females had significantly more total egg chambers per ovariole compared to YLB females (Table S3). Pairwise comparisons with Tukey’s adjustment indicated that this difference was particularly pronounced in mated conditions (9.01% more in mated BL females, Table S4). The proportion of pre-vitellogenic egg chambers showed no significant effect from selection regime, mating status, or their interaction (Table 1, Figure 4c). For vitellogenic egg chambers, the main effects of selection regime and the effect of selection regime × mating status interaction were not significant (Table 2). However, mating status had a significant effect (Table 2, Figure 4c), wherein regardless of selection regime, mated females were found to have 55.02 % more vitellogenic egg chambers than virgin females (Table S4, Figure 4c). Analysis of the proportion of post-vitellogenic egg chambers indicated significant effects of both selection regime and mating status (Table 2, Figure 4c). YLB females were found to possess ∼20% more post-vitellogenic stages than that in the BL females (Table S4). Virgin females were found to have a higher number of mature oocytes regardless of selection regime (Table S4). The effect of interaction between selection regime and mating status on the proportion of post-vitellogenic egg chambers was marginally non-significant (Table 2).

**Figure 4.**
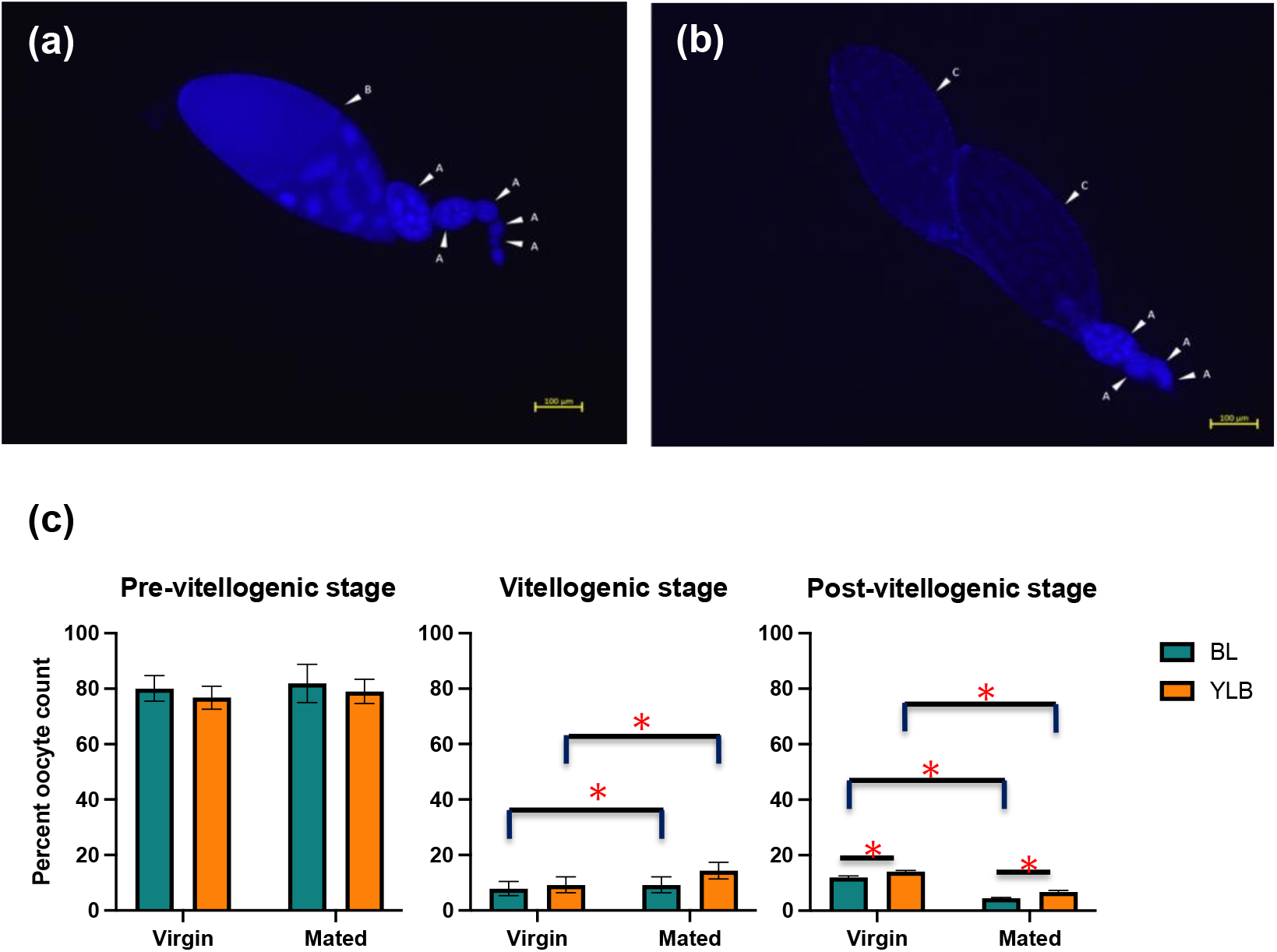
Important results from Experiment 6. (a) a typical DAPI stained image of an ovariole from a BL female, (b) a typical DAPI stained image of an ovariole from a YLB female. The arrowheads points to the different stages of egg chambers. A: Pre-vitellogenic, B: vitellogenic, and C: post-vitellogenic. (c): Percentage of egg chambers at different developmental stages from Experiment 6. The percentage of pre-vitellogenic, vitellogenic, and post-vitellogenic egg chambers was quantified for virgin and mated females. The cyan bar represents data from BL (control regime) females, while the orange bar represents data from YLB (experimental regime) females. The error bar represents standard error of means. Statistically significant differences, as indicated by multiple comparisons, are indicated with an asterisk (*).

## Discussion

Reproduction is a protein-intensive process and hence, most animals need persistent access to proteins to reproduce (Skorupa et al., 2008). We found that as a result of adaptation to chronic protein deprivation experimental populations (YLB) evolved significantly higher larval feeding rate, and an associated increase in whole-body weight and protein content at the time of eclosion (emergence of an adult from puparium). Thus, the results indicate the possibility of evolution of nutrient specific resource acquisition and/or assimilation at the larval stage. Interestingly, YLB females were also found to have a higher body mass and absolute lipid content in the fitness window – indicating the evolution of adult stage resource acquisition and/or differential utilization and storage as well. Our results further suggest changes in ovarian size (ovariole count) and function (production of mature oocytes) aiding the attainment of a higher rate of egg production. The YLB females maintained a higher number of ovarioles and a greater number of mature (post-vitellogenic) oocytes per ovariole. These results shed light on the proximate mechanisms underlying the attainment of a higher early-life fecundity, and indicate the putative role of genetic variance and covariance between juvenile and adult fitness traits pertinent to such evolution.

### Resource acquisition and storage

In holometabolous insects, early-life reproduction is reliant on larval resource acquisition and storage (Telang & Wells, 2004; Aguila et al., 2013). In *D. melanogaster*, ovarian development begins during metamorphosis (Bodenstein, 1950) and hence, early-life ovarian output could rely primarily on larval nutrition. Unsurprisingly, YLBs were found to have an increased early-third instar feeding rate, the most prolific feeding stage (Li et al., 2008; Wang et al., 2023). Evolution of feeding rate under a variety of life history selection is fairly well-known in the literature (Joshi & Mueller, 1988; Rajamani et al., 2006; Mueller & Barter, 2015; Bitner et al., 2021). However, such change in feeding behaviour is expected to result in an increase in overall body mass at eclosion (Bierbaum et al., 1989; Klepsatel et al., 2018). This trend was not observed in our YLB females. Instead, only (whole-body) protein content seemed to have increased, at least at the time of eclosion. To the best of our knowledge, this is the first report of a macronutrient-nutrient specific effect of increased feeding rate. Nonetheless, since protein is known to be a limiting macronutrient for egg production, such an observation is consistent with the *macronutrient content theory* mentioned earlier. The observed increase in protein content on eclosion is expected to have allowed the attainment of elevated early-life fecundity, especially under a protein restricted ecology, assuming that at least part of this protein reserve is eventually allocated to egg production. Curiously, increased whole body protein content was not observed at the fitness window (i.e., 4-5 days post-eclosion). Given that Bradford assay is less sensitive to conjugated proteins in the body (Giraudi et al., 1997), the observed results could indicate reproductive allocation of stored protein, as much of the protein molecules in the developing oocyte, especially yolk protein such as, vitellogenin, is usually found in conjugated forms such as glycolipid and phosphoprotein (Raikhel & Dhadialla, 1992). Thus, the observed results on protein content could simply indicate reproductive allocation of stored protein.

Apart from juvenile phase resource acquisition, adult foraging can significantly contribute to biomass accumulation that can boost reproductive output (Partridge et al., 1999). Our results on whole body dry weight corroborated this theory and showed that body mass in females increased from eclosion to the fitness window, i.e., during 4-5 days of adult life. Interestingly, the evolved YLB females showed a greater increase compared to the control females, indicating that the evolved females were putting on more biomass during the adult phase until the reproductive (fitness) window. Such biomass accumulation can either be specific to the reproductive system, i.e., in reproductive structures – especially increased ovary size (Simmons & Bradley, 1997), or be more generic long-term storage, especially lipid. The latter would naturally make individuals more resistant to starvation (Chippindale, Chu & Rose, 1996; Rion & Kawecki, 2007; Rehman & Varghese, 2021), while the former would allow females to be more fecund.

### Allocation of acquired resource to ovaries

Two lines of evidence uphold the theory of reproductive biomass accumulation. First, YLB females were found to have significantly higher total body lipid content at the time of their reproductive window, without a correspondingly higher resistance to starvation. This implied that stored lipid is not available to general somatic metabolism, and perhaps was locked in reproductive tissues. Importantly, *Drosophila* ovaries are known to contain ∼15% of the whole-body lipid content (Lease & Wolf, 2011; Colinet & Renault, 2014). It is important to note that the observed increase in lipid content among the evolved females corresponds with their increase in whole-body biomass. This suggests that changes in resource acquisition and/or utilization during the adult phase, which are reflected in biomass accumulation, primarily influenced lipid storage. Secondly, we found ovariole count, a proxy for ovary size (Mendes & Mirth, 2016; Sarikaya et al., 2019), to be higher in YLB females. Since a substantial part of ovarian biomass is known to be lipid, it is not unreasonable to suggest that the increase in lipid content discussed earlier corresponds to increase in ovary size, at least partly. Thus, our results appear to suggest an evolution of larval and adult macronutrient specific resource assimilation, but a preferential allocation of it to the reproductive system. The evolution of such preferential allocation of metabolic resources is only beginning to emerge (also see Vijendravarma, 2018). It is important to understand how this can translate into changes in egg production.

### Ovarian function

Notwithstanding the limited information in the literature (Hodin, 2009; Sarikaya et al., 2012; Mendes & Mirth, 2016), our results suggest plasticity in ovariole count in females, such that the ovariole count in mated females was significantly less than that in the virgin females (but see Carlson et al., 1998). Putatively, programmed cell death, which is commonly observed in insect ovaries, including *Drosophila* sp., can potentially modulate ovariole number (Pritchett et al., 2009; Almeida & Suesdek, 2017; Ronai et al., 2016, 2017). In adult fruit flies, protein restriction leads to increased cell death in the germaria and the mid-stages of oogenesis (Drummond-Barbosa et al., 2001; McCall, 2004). In face of nutritional stress, cell deaths are induced in these regions which act as checkpoints determining whether egg chambers should proceed to the vitellogenesis process (Drummond-Barbosa et al., 2001). Mating is known to promote increased oogenesis (Soller et al., 1997; Avila et al., 2011; McDonough-Goldstein et al., 2021). It is not unreasonable to suggest that rapid increase in oogenesis quickly becomes unsustainable, especially under sub-optimal diets such as, protein restriction similar to our assay condition -triggering programmed cell death of developing egg chambers and/or ovarioles. This could indeed act as an effective checkpoint, and regulation on reproductive investment (Moore & Attisano, 2011; Adler & Bonduriansky, 2014).

Interestingly, we did not detect this difference in ovariole count across the two mating statuses in evolved (YLB) females. This could have been achieved either by evolving a higher threshold for the apoptotic control to set in, and/or by making more metabolic resources (e.g., protein, lipid etc.) available to the reproductive tissues. While the latter seems likely, the former cannot be ruled out at this point.

Interestingly, our results on oocyte developmental stages suggested that although the ovaries of the females of the control population had significantly more total egg chambers, the evolved females produced a higher number of post-vitellogenic egg chambers. There are at least three ways in which this could have been achieved by the evolved females – (a) increased rate of oocyte development and oviposition, (b) reduced oocyte arrest at the pre-vitellogenic stages, or (c) reduced apoptosis at the checkpoints (Drummond-Barbosa, 2001). These three possibilities are not mutually exclusive. Further investigations are needed to test these theories. Irrespective of the regime, ovaries of mated females were found to possess fewer mature egg chambers probably because mating stimulated oviposition (Carlson et al., 1998).

### Evolutionary genetics of nutrition-dependent adaptation

Response in experimental evolution usually depends on standing genetic variation (Teotónio et al. 2009, Kawecki et al. 2012). Hence, our results directly indicate the presence of a substantial standing genetic variation in resource acquisition, storage, and allocation traits, in line with the existing literature (Joshi & Mueller, 1996; Hoffmann & Harshman, 1999; Rion & Kawecki, 2007). Further, though biomass at eclosion seems to be genetically correlated with macronutrient content (Chippindale et al. 1996, Kristensen et al. 2011, Enriquez et al. 2022), it is evident that protein and lipid content at eclosion are, at least, partly genetically uncorrelated. Whether these results were due to changes in protein uptake during larval stage, or metabolic conversion of other macronutrients to protein, should be a topic of future investigations. Dasgupta et al. (2022) posed several plausible scenarios in which females can maximize early-life egg output without a corresponding trade-off with lifespan. We now know that, not only reduction in late-life fecundity (Dasgupta et al., 2022), increased resource availability to reproduction, which culminated in increased ovarian function was the mechanistic basis of the observed nutrient dependent adaptation.

Interestingly, we showed mating-dependent plasticity in ovariole number and rate of production of mature stages of oocyte, and the evolution of such plasticity. To the best of our knowledge this is the first report of such evolution of female reproductive physiology. Evolution of phenotypic plasticity itself is a fairly underexplored area (but see Maggu et al., 2020). Whether such evolution is fundamentally reliant on the evolution of control elements of ovarian function remains to be explored. Nonetheless, our results are indicative of plausible involvement of some signaling pathway that connects nutrient sensing and ovarian function (Durdevic & Ephrussi, 2019), an area yet to be fully explored.

## Author contributions

BN conceived the idea, designed the experiment, analysed and interpreted the data, edited the manuscript, and secured funding. PD and AK designed and performed the experiments, analysed data. PD prepared the first draft of the manuscript. RSP, PNP, and KR performed various assays and collected data.

## Acknowledgements

We are thankful to Tanya Verma and Subhasish Halder for their help during the experiments and population maintenance. The experiments reported here was supported by the Core Research Grant (file no. CRG/2019/002460) from Science and Engineering Research Board, Department of Science and Technology, Government of India. PD thanks Council for Scientific and Industrial Research, Government of India for financial support in the form of Junior and Senior Research Fellowship.

## References

Adler, M. I., & Bonduriansky, R. (2014). Why do the well-fed appear to die young? A new evolutionary hypothesis for the effect of dietary restriction on lifespan. Bioessays, 36, 439–450.

Aguila, J. R., Hoshizaki, D. K., & Gibbs, A. G. (2013). Contribution of larval nutrition to adult reproduction in Drosophila melanogaster. Journal of Experimental Biology, 216(3), 399–406.

Ahmad, M., Keebaugh, E. S., Tariq, M., & Ja, W. W. (2018). Evolutionary responses of Drosophila melanogaster under chronic malnutrition. Frontiers in ecology and evolution, 6, 47.

Almeida, F., & Suesdek, L. (2017). Effects of Wolbachia on ovarian apoptosis in Culex quinquefasciatus (Say, 1823) during the previtellogenic and vitellogenic periods. Parasites and Vectors, 10(1), 1–8.

Avila, F. W., Sirot, L. K., Laflamme, B. A., Rubinstein, C. D., & Wolfner, M. F. (2011). Insect seminal fluid proteins: Identification and function. Annual Review of Entomology, 56, 21–40.

Ballard, J. W. O., Melvin, R. G., & Simpson, S. J. (2008). Starvation resistance is positively correlated with body lipid proportion in five wild caught Drosophila simulans populations. Journal of insect physiology, 54(9), 1371–1376.

Bates, D., Mächler, M., Bolker, B., & Walker, S. (2014). Fitting linear mixed-effects models using lme4. arXiv preprint arXiv:1406.5823.

Bierbaum, T. J., Mueller, L. D., & Ayala, F. J. (1989). Density-Dependent Evolution of Life-History Traits in Drosophila melanogaster. Evolution, 43(2), 382.

Bitner, K., Rutledge, G. A., Kezos, J. N., & Mueller, L. D. (2021). The effects of adaptation to urea on feeding rates and growth in Drosophila larvae. Ecology and evolution, 11(14), 9516–9529.

Bodenstein, D. (1950). The postembryonic development of Drosophila. Biology of Drosophila, 275–367.

Boggs, C. L. (1992). Resource allocation: exploring connections between foraging and life history. Functional Ecology, 6(5), 508–518.

Bonduriansky, R., Maklakov, A., Zajitschek, F., & Brooks, R. (2008). Sexual selection, sexual conflict and the evolution of ageing and life span. Functional Ecology, 22(3), 443–453.

Bradford, M. M. (1976). A rapid and sensitive method for the quantitation of microgram quantities of protein utilizing the principle of protein-dye binding. Analytical biochemistry, 72(1-2), 248–254.

Carey, J. R., Harshman, L. G., Liedo, P., Müller, H. G., Wang, J. L., & Zhang, Z. (2008). Longevity–fertility trade-offs in the tephritid fruit fly, Anastrepha ludens, across dietary-restriction gradients. Aging cell, 7(4), 470–477.

Carlson, K. A., Nusbaum, T. J., Rose, M. R., & Harshman, L. G. (1998). Oocyte maturation and ovariole number in lines of Drosophila melanogaster selected for postponed senescence. Functional Ecology, 12(4), 514–520.

Cavigliasso, F., Dupuis, C., Savary, L., Spangenberg, J. E., & Kawecki, T. J. (2020). Experimental evolution of post-ingestive nutritional compensation in response to a nutrient-poor diet. Proceedings of the Royal Society B, 287(1940), 20202684.

Chippindale, A. K., Chu, T. J. F., & Rose, M. R. (1996). Complex trade-offs and the evolution of starvation resistance in Drosophila melanogaster. Evolution, 50(2), 753–766.

Chippindale, A. K., Leroi, A. M., Kim, S. B., & Rose, M. R. (1993). Phenotypic plasticity and selection in Drosophila life-history evolution. I. Nutrition and the cost of reproduction. Journal of Evolutionary Biology, 6(2), 171–193.

Churchill, E. R., Dytham, C., & Thom, M. D. (2019). Differing effects of age and starvation on reproductive performance in Drosophila melanogaster. Scientific reports, 9(1), 2167.

Colinet, H., & Renault, D. (2014). Dietary live yeast alters metabolic profiles, protein biosynthesis and thermal stress tolerance of Drosophila melanogaster. Comparative Biochemistry and Physiology - A Molecular and Integrative Physiology, 170, 6–14.

Coyne, J. A., Rux, J., & David, J. R. (1991). Genetics of morphological differences and hybrid sterility between Drosophila sechellia and its relatives. Genetics Research, 57(2), 113–122.

Dasgupta, P., Halder, S., Dari, D., Nabeel, P., Vajja, S. S., & Nandy, B. (2022). Evolution of a novel female reproductive strategy in Drosophila melanogaster populations subjected to long-term protein restriction. Evolution, 76(8), 1836–1848.

Desprez, M., Gimenez, O., McMahon, C. R., Hindell, M. A., & Harcourt, R. G. (2018). Optimizing lifetime reproductive output: Intermittent breeding as a tactic for females in a long-lived, multiparous mammal. Journal of Animal Ecology, 87(1), 199–211.

Drummond-Barbosa, D., & Spradling, A. C. (2001). Stem cells and their progeny respond to nutritional changes during Drosophila oogenesis. Developmental biology, 231(1), 265–278.

Durdevic, Z., & Ephrussi, A. (2019). Germ cell lineage homeostasis in Drosophila requires the Vasa RNA helicase. Genetics, 213(3), 911–922.

Ellison, P. T. (2003). Energetics and reproductive effort. American Journal of Human Biology, 15(3), 342–351.

Enriquez, T., Lievens, V., Nieberding, C. M., & Visser, B. (2022). Pupal size as a proxy for fat content in laboratory-reared and field-collected Drosophila species. Scientific Reports, 12(1), 12855.

Fontana, L., & Partridge, L. (2015). Promoting health and longevity through diet: From model organisms to humans. Cell, 161(1), 106–118.

Foray, V., Pelisson, P. F., Bel-Venner, M. C., Desouhant, E., Venner, S., Menu, F., … & Rey, B. (2012). A handbook for uncovering the complete energetic budget in insects: the van Handel’s method (1985) revisited. Physiological Entomology, 37(3), 295–302.

Gatto, M., Matessi, C., & Slobodkin, L. B. (1989). Physiological profiles and demographic rates in relation to food quantity and predictability: an optimization approach. Evolutionary Ecology, 3, 1–30.

Giraudi, G., Baggiani, C., & Giovannoli, C. (1997). Inaccuracy of the Bradford method for the determination of protein concentration in steroid-horseradish peroxidase conjugates. Analytica chimica acta, 337(1), 93–97.

Hodin, J. (2009). She shapes events as they come: Plasticity in female insect reproduction. Phenotypic Plasticity of Insects : Mechanisms and Consequences, 99.

Hoffmann, A. A., & Harshman, L. G. (1999). Desiccation and starvation resistance in Drosophila: Patterns of variation at the species, population and intrapopulation levels. Heredity, 83(6), 637–643.

Inness, C. L., & Metcalfe, N. B. (2008). The impact of dietary restriction, intermittent feeding and compensatory growth on reproductive investment and lifespan in a short-lived fish. Proceedings of the Royal Society B: Biological Sciences, 275(1644), 1703–1708.

Jervis, M. A., Ellers, J., & Harvey, J. A. (2008). Resource acquisition, allocation, and utilization in parasitoid reproductive strategies. Annu. Rev. Entomol., 53(1), 361–385.

Jia, D., Xu, Q., Xie, Q., Mio, W., & Deng, W. M. (2016). Automatic stage identification of Drosophila egg chamber based on DAPI images. Scientific reports, 6(1), 18850.

Joshi, A., & Mueller, L. D. (1988). Evolution of higher feeding rate in Drosophila due to density-dependent natural selection. Evolution, 42, 1090– 1093.

Joshi, A., & Mueller, L. D. (1996). Density-dependent natural selection in Drosophila: Trade-offs between larval food acquisition and utilization. Evolutionary Ecology, 10(5), 463–474.

Kawecki, T. J., Lenski, R. E., Ebert, D., Hollis, B., Olivieri, I., & Whitlock, M. C. (2012). Experimental evolution. Trends in Ecology and Evolution, 27(10), 547–560.

Kirkwood, T. B. (1977). Evolution of ageing. Nature, 270(5635), 301–304.

Klepsatel, P., Procházka, E., & Gáliková, M. (2018). Crowding of Drosophila larvae affects lifespan and other life-history traits via reduced availability of dietary yeast. Experimental Gerontology, 110, 298–308.

Kolss, M., Vijendravarma, R. K., Schwaller, G., & Kawecki, T. J. (2009). Life-history consequences of adaptation to larval nutritional stress in Drosophila. Evolution, 63(9), 2389– 2401.

Kristensen, T. N., Overgaard, J., Loeschcke, V., & Mayntz, D. (2011). Dietary protein content affects evolution for body size, body fat and viability in Drosophila melanogaster. Biology letters, 7(2), 269–272.

Kristensen, T. N., Ketola, T., & Kronholm, I. (2020). Adaptation to environmental stress at different timescales. Annals of the New York Academy of Sciences, 1476(1), 5–12.

Kuznetsova, A., Brockhoff, P. B., & Christensen, R. H. (2017). lmerTest package: tests in linear mixed effects models. Journal of statistical software, 82, 1–26.

Lease, H. M., & Wolf, B. O. (2011). Lipid content of terrestrial arthropods in relation to body size, phylogeny, ontogeny and sex. Physiological Entomology, 36(1), 29–38.

Lee, K. P., & Jang, T. (2014). Exploring the nutritional basis of starvation resistance in Drosophila melanogaster. Functional Ecology, 28(5), 1144–1155.

Lenth, R., Singmann, H., Love, J., Buerkner, P., & Herve, M. (2018). Emmeans: Estimated marginal means, aka least-squares means. R package version, 1, 3.

Li, H. M., Buczkowski, G., Mittapalli, O., Xie, J., Wu, J., Westerman, R., … & Pittendrigh, B. R. (2008). Transcriptomic profiles of Drosophila melanogaster third instar larval midgut and responses to oxidative stress. Insect molecular biology, 17(4), 325–339.

Lobell, A. S., Kaspari, R. R., Serrano Negron, Y. L., & Harbison, S. T. (2017). The genetic architecture of Ovariole number in Drosophila melanogaster: Genes with major, quantitative, and pleiotropic effects. G3: Genes, Genomes, Genetics, 7(7), 2391–2403.

Long, T. A. F., Pischedda, A., Nichols, R. V., & Rice, W. R. (2010). The timing of mating influences reproductive success in Drosophila melanogaster: implications for sexual conflict. Journal of Evolutionary Biology, 23(5), 1024–1032.

Maggu, K., Ahlawat, N., Geeta Arun, M., Meena, A., & Prasad, N. G. (2021). Divergence of responses to variable socio-sexual environments in laboratory populations of Drosophila melanogaster evolving under altered operational sex ratios. Evolution, 75(2), 414–426.

Magwere, T., Chapman, T., & Partridge, L. (2004). Sex differences in the effect of dietary restriction on life span and mortality rates in female and male Drosophila melanogaster. The Journals of Gerontology Series A: Biological Sciences and Medical Sciences, 59(1), B3–B9.

Mair, W., Sgrò, C. M., Johnson, A. P., Chapman, T., & Partridge, L. (2004). Lifespan extension by dietary restriction in female Drosophila melanogaster is not caused by a reduction in vitellogenesis or ovarian activity. Experimental gerontology, 39(7), 1011–1019.

Mair, W., Piper, M. D. W., & Partridge, L. (2005). Calories do not explain extension of life span by dietary restriction in Drosophila. PLoS Biology, 3(7), 1305–1311.

Marron, M. T., Markow, T. A., Kain, K. J., & Gibbs, A. G. (2003). Effects of starvation and desiccation on energy metabolism in desert and mesic Drosophila. Journal of Insect Physiology, 49(3), 261–270.

McCall, K. (2004). Eggs over easy: Cell death in the Drosophila ovary. Developmental Biology, 274(1), 3–14.

McDonough-Goldstein, C. E., Pitnick, S., & Dorus, S. (2021). Drosophila oocyte proteome composition covaries with female mating status. Scientific reports, 11(1), 3142.

Mendes, C. C., & Mirth, C. K. (2016). Stage-specific plasticity in ovary size is regulated by insulin/insulin-like growth factor and ecdysone signaling in Drosophila. Genetics, 202(2), 703– 719.

Moatt, J. P., Nakagawa, S., Lagisz, M., & Walling, C. A. (2016). The effect of dietary restriction on reproduction: a meta-analytic perspective. BMC Evolutionary Biology, 16, 1–9.

Moatt, J. P., Savola, E., Regan, J. C., Nussey, D. H., & Walling, C. A. (2020). Lifespan extension via dietary restriction: time to reconsider the evolutionary mechanisms?. BioEssays, 42, 1900241.

Moore, P. J., & Attisano, A. (2011). Oosorption in response to poor food: Complexity in the trade-off between reproduction and survival. Ecology and Evolution, 1(1), 37–45.

Mueller, L. D., & Barter, T. T. (2015). A model of the evolution of larval feeding rate in Drosophila driven by conflicting energy demands. Genetica, 143, 93–100.

Nakagawa, S., Lagisz, M., Hector, K. L., & Spencer, H. G. (2012). Comparative and meta-analytic insights into life extension via dietary restriction. Aging Cell, 11(3), 401–409.

Nandy, B., Joshi, A., Ali, Z. S., Sen, S., & Prasad, N. G. (2012). Degree of adaptive male mate choice is positively correlated with female quality variance. Scientific Reports, 2, 1–8.

Partridge, L., Langelan, R., Fowler, K., Zwaan, B., & French, V. (1999). Correlated responses to selection on body size in Drosophila melanogaster. Genetics Research, 74(1), 43–54.

Partridge, L., Piper, M. D. W., & Mair, W. (2005). Dietary restriction in Drosophila. Mechanisms of Ageing and Development, 126(9 SPEC. ISS.), 938–950.

Pianka, E. R. (1976). Natural selection of optimal reproductive tactics. American Zoologist, 16(4), 775–784.

Piper, M. D., Mair, W., & Partridge, L. (2005). Counting the calories: the role of specific nutrients in extension of life span by food restriction. The Journals of Gerontology Series A: Biological Sciences and Medical Sciences, 60, 549–555.

Pritchett, T. L., Tanner, E. A., & McCall, K. (2009). Cracking open cell death in the Drosophila ovary. Apoptosis, 14(8), 969–979.

Raikhel, A. S., & Dhadialla, T. S. (1992). Accumulation of yolk proteins in insect oocytes.

Rajamani, M., Raghavendra, N., Prasad, N. G., Archana, N., Joshi, A., & Shakarad, M. (2006). Reduced larval feeding rate is a strong evolutionary correlate of rapid development in Drosophila melanogaster. Journal of Genetics, 85(3), 209–212.

Rehman, N., & Varghese, J. (2021). Larval nutrition influences adult fat stores and starvation resistance in Drosophila. PLoS One, 16(2), e0247175.

Rion, S., & Kawecki, T. J. (2007). Evolutionary biology of starvation resistance: What we have learned from Drosophila. Journal of Evolutionary Biology, 20(5), 1655–1664.

Roff, D. (Ed.). (1993). Evolution of life histories: theory and analysis. Springer Science & Business Media.

Ronai, I., Allsopp, M. H., Tan, K., Dong, S., Liu, X., Vergoz, V., & Oldroyd, B. P. (2017). The dynamic association between ovariole loss and sterility in adult honeybee workers. Proceedings of the Royal Society B: Biological Sciences, 284(1851).

Ronai, I., Oldroyd, B. P., & Vergoz, V. (2016). Queen pheromone regulates programmed cell death in the honey bee worker ovary. Insect Molecular Biology, 25(5), 646–652.

Sarikaya, D. P., Belay, A. A., Ahuja, A., Dorta, A., Green, D. A., & Extavour, C. G. (2012). The roles of cell size and cell number in determining ovariole number in Drosophila. Developmental Biology, 363(1), 279–289.

Sarikaya, D. P., Church, S. H., Lagomarsino, L. P., Magnacca, K. N., Montgomery, S. L., Price, D. K., … Extavour, C. G. (2019). Reproductive Capacity Evolves in Response to Ecology through Common Changes in Cell Number in Hawaiian Drosophila. Current Biology, 29(11), 1877–1884.

Schwasinger-schmidt, T. E., Kachman, S. D., & Harshman, L. G. (2012). Evolution of starvation resistance in Drosophila melanogaster: measurement of direct and correlated responses to artificial selection. Journal of evolutionary biology, 25(2), 378–387.

Shanley, D. P., & Kirkwood, T. B. (2000). Calorie restriction and aging: A life-history analysis. Evolution, 54(3), 740–750.

Shu, R., Uy, L., & Wong, A. C. N. (2022). Nutritional phenotype underlines the performance trade-offs of Drosophila suzukii on different fruit diets. Current Research in Insect Science, 2, 100026.

Simmons, F. H., & Bradley, T. J. (1997). An analysis of resource allocation in response to dietary yeast in Drosophila melanogaster. Journal of insect physiology, 43(8), 779–788.

Skorupa, D. A., Dervisefendic, A., Zwiener, J., & Pletcher, S. D. (2008). Dietary composition specifies consumption, obesity, and lifespan in Drosophila melanogaster. Aging cell, 7(4), 478– 490.

Soller, M., Bownes, M., & Kubli, E. (1997). Mating and sex peptide stimulate the accumulation of yolk in oocytes of Droosophila melanogaster. European Journal of Biochemistry, 243(3), 732–738.

Stearns, S. C. (1989). Trade-offs in life-history evolution. Functional ecology, 3(3), 259–268.

Stearns, S. C. (1992). The evolution of life histories (Vol. 249, p. xii). Oxford: Oxford university press.

Stearns, S. C. (2000). Life history evolution: successes, limitations, and prospects. Naturwissenschaften, 87(11), 476–486.

Telang, A., & Wells, M. A. (2004). The effect of larval and adult nutrition on successful autogenous egg production by a mosquito. Journal of Insect Physiology, 50(7), 677–685.

Teleman, A. A. (2010). Molecular mechanisms of metabolic regulation by insulin in Drosophila. Biochemical Journal, 425(1), 13–26.

Teotónio, H., Chelo, I. M., Bradić, M., Rose, M. R., & Long, A. D. (2009). Experimental evolution reveals natural selection on standing genetic variation. Nature genetics, 41(2), 251–257.

Vijendravarma, R. K. (2018). Experimental evolution demonstrates evolvability of preferential nutrient allocation to competing traits in response to chronic malnutrition. Journal of Evolutionary Biology, 31(11), 1743–1749.

Wang, L., Wei, D. D., Wang, G. Q., Huang, H. Q., & Wang, J. J. (2023). High-sucrose diet exposure on larvae contributes to adult fecundity and insecticide tolerance in the oriental fruit fly, Bactrocera dorsalis (Hendel). Insects, 14(5), 407.

Wayne, M. L., Soundararajan, U., & Harshman, L. G. (2006). Environmental stress and reproduction in Drosophila melanogaster: Starvation resistance, ovariole numbers and early age egg production. BMC Evolutionary Biology, 6, 1–10.

Williams, G. C. (1957). Pleiotropy, natural selection, and the evolution of senescence. Evolution, 11, 398–411.

Zajitschek, F., Georgolopoulos, G., Vourlou, A., Ericsson, M., Zajitschek, S. R. K., Friberg, U., & Maklakov, A. A. (2019). Evolution under dietary restriction decouples survival from fecundity in Drosophila melanogaster females. Journals of Gerontology - Series A Biological Sciences and Medical Sciences, 74(10), 1542–1548.

Zajitschek, F., Zajitschek, S. R., Canton, C., Georgolopoulos, G., Friberg, U., & Maklakov, A. A. (2016). Evolution under dietary restriction increases male reproductive performance without survival cost. Proceedings of the Royal Society B: Biological Sciences, 283(1825), 20152726.

Zimmerman, J. A., Malloy, V., Krajcik, R., & Orentreich, N. (2003). Nutritional control of aging. Experimental gerontology, 38, 47–52.

