## Supplementary information for "Adaptation to yeast restriction through the evolution of resource acquisition, macronutrient reserve, and ovarian function"

**S1. Bradford assay and the measurement of protein content:**

The whole-body protein content was quantified for the group of flies collected at two time points – i.e., on eclosion and at the fitness window. Bradford method was used for this purpose (Bradford 1976, Foray et al. 2012). For the preparation of the standard curve, Bovine Serum Albumin solutions of concentration 1000 µg/ml, 500 µg/ml and 250 µg/ml were prepared using double-distilled water. Double-distilled water was used as a “blank”. Each set of three females, kept in the vials, were transferred to 1.5ml centrifuge tubes with 360µl of lysis buffer (100mM KH_2_PO_4_, 1mM Ethylene-diamine-tetraacetate, 1mM Dichloro-diphenyl-trichloroethane, pH = 7.4). The samples were homogenized properly using a micro pestle, and centrifuged at 2000 rpm for 13 minutes at 4℃. In a 96-well plate, for each sample as well as the standard solution, 5µl of supernatant was suspended in three wells, which meant each sample had three technical replicates. Thereafter, 250µl of Bradford Reagent (*Sigma-Aldrich*) was added in each well. After 15-20 minutes of incubation at room temperature (~ 25-30℃), absorbance of the samples in the plate was recorded using a spectrophotometer (*Epoch 2, BioTek Instruments*).

Data processing

All three absorbance values of the technical replicates for each sample were averaged and subtracted by the average absorbance value of the blank. Obtained values of absorbance for each sample were used for further calculation. A standard curve was prepared by fitting the standard concentration values of Bovine Serum Albumin solutions (i.e., 1000 µg/ml, 500 µg/ml and 250 µg/ml) on the corresponding absorbance values. By fitting the observed absorbance values to the standard curve, protein concentration of the experimental samples was obtained. For the statistical analyses, whole body protein content was computed from these protein concentration values using a method described in the supplementary information (refer to the Supplementary Information: S2). For the final analysis, after the exclusion of outliers, values from 10-12 samples were considered for each population.

**S2. Detailed calculation for protein content data processing**

The calculation for quantifying the whole-body protein content has been discussed here in details using a trial dataset. The absorbance values for the standard and the samples, so obtained by Bradford assay have been mentioned here:

| **Concentration** | **Absorbance** |
| --- | --- |
| 1000 µg/ml BSA | 0.34 |
| 500 µg/ml BSA | 0.25 |
| 250 µg/ml BSA | 0.12 |
| Control (DDW) | 0 |
| Sample | 0.17967 |

The concentrations of BSA and their corresponding absorbance values are plotted in X and Y axis respectively. The standard curve equation was:

y = 0.0299 + 0.0003x; x = (y - 0.0299)/0.0003

By substituting y with the absorbance value of the sample in the equation, we obtain its concentration as x.

x = (0.17967-0.0299)/0.0003; x = 499.22µg/ml

According to the protocol, 3 flies were homogenized in 360µl of lysis buffer. Hence, we calculated the amount of protein in 360µl of the sample, i.e. (499.22/1000)$\times$360 = 179.72µg of protein in 360µl of sample.

In the trial, the mean dry mass of the females in a population was measured to be 0.387mg. Since, 360µl of a sample contains 3 flies homogenized in it, the protein content value is divided by 3 times the mean dry mass to obtain the normalized protein content. 79.72/ (3$\times$0.387)= 154.80µg/mg

| **Contrasts** | **p value** |
| --- | --- |
| BL FW- YLB FW | **<0.01** |
| BL OE - YLB OE | 0.15 |
| BL OE-BL FW | **<0.01** |
| YLB OE- YLB FW | **<0.01** |

Table S1: Outcome of the post-hoc analysis of dry body mass comparison between BL and YLB females on eclosion (OE) and in fitness window (FW). Tukey’s adjusted p-values were obtained by pairwise comparisons using *emmeans* package in R version 4.0.3. Statistically significant p-values are mentioned in bold face.

| **Contrasts** | **p value** |
| --- | --- |
| BL mated- BL virgin | **<0.01** |
| YLB mated- YLB virgin | 0.88 |
| BL virgin-YLB virgin | 0.15 |
| BL mated- YLB mated | **0.03** |

Table S2: Outcome of the post-hoc analysis of mean ovariole count comparison between BL and YLB females under virgin and mated conditions. Tukey’s adjusted p-values were obtained by pairwise comparisons using *emmeans* package in R version 4.0.3. Statistically significant p-values are mentioned in bold face.

| **Contrasts** | **p-value** |
| --- | --- |
| BL mated- BL virgin | 0.44 |
| YLB mated-YLB virgin | **<0.01** |
| BL virgin- YLB virgin | 0.4 |
| BL mated- YLB mated | **<0.01** |

Table S3: Outcome of the post-hoc analysis of total developing oocyte count comparison between BL and YLB females under virgin and mated conditions. Tukey’s adjusted p-values were obtained by pairwise comparisons using *emmeans* package in R version 4.0.3. Statistically significant p-values are mentioned in bold face.

| **Sl. No.** | **Traits** | **Time point/ Mating status** | **Sel.reg.** | **Mean** | **Std Err** |
| --- | --- | --- | --- | --- | --- |
| **1** | **Larval feeding rate** | N/A | BL | 145.55 | 2.23 |
|  |  |  | YLB | 163.78 | 2.22 |
| **2** | **Protein content (μg/mg)** | On eclosion | BL | 174.72 | 4.19 |
|  |  |  | YLB | 201.78 | 4.2 |
|  |  | Fitness window | BL | 123.49 | 4.06 |
|  |  |  | YLB | 130.19 | 4.06 |
| **3** | **Absolute lipid mass (μg)** | On eclosion | BL | 54.32 | 1.59 |
|  |  |  | YLB | 46.26 | 1.59 |
|  |  | Fitness window | BL | 63.44 | 1.75 |
|  |  |  | YLB | 73.72 | 1.69 |
| **4** | **Relative lipid content (%)** | On eclosion | BL | 15.37 | 0.39 |
|  |  |  | YLB | 13.01 | 0.39 |
|  |  | Fitness window | BL | 15.68 | 0.39 |
|  |  |  | YLB | 17.51 | 0.37 |
| **5** | **Dry body mass (μg)** | On eclosion | BL | 354.82 | 2.04 |
|  |  |  | YLB | 361.47 | 1.98 |
|  |  | Fitness window | BL | 398.25 | 2.12 |
|  |  |  | YLB | 420.17 | 2.02 |
| **6** | **Starvation resistance (in hours)** | N/A | BL | 96.32 | 0.89 |
|  |  |  | YLB | 99.34 | 0.92 |
| **7** | **Mean ovariole count** | Virgin | BL | 20.01 | 0.23 |
|  |  |  | YLB | 19.45 | 0.22 |
|  |  | Mated | BL | 18.63 | 0.23 |
|  |  |  | YLB | 19.51 | 0.23 |
| **8** | **Thorax length (in µm)** | Virgin | BL | 1059.16 | 4.38 |
|  |  |  | YLB | 1052.59 | 4.42 |
|  |  | Mated | BL | 1063.47 | 4.35 |
|  |  |  | YLB | 1053.85 | 4.17 |
| **9** | **Total oocyte stages per unit ovariole** | Virgin | BL | 6.58 | 0.08 |
|  |  |  | YLB | 6.48 | 0.35 |
|  |  | Mated | BL | 6.72 | 0.88 |
|  |  |  | YLB | 6.12 | 0.46 |
| **10** | **Pre-vitellogenic egg chambers** | Virgin | BL | 100.85 | 2.29 |
|  |  |  | YLB | 92.48 | 2.18 |
|  |  | Mated | BL | 94.38 | 2.3 |
|  |  |  | YLB | 91.02 | 2.26 |
| **11** | **Vitellogenic egg chambers** | Virgin | BL | 10.06 | 0.57 |
|  |  |  | YLB | 11.04 | 0.51 |
|  |  | Mated | BL | 16.23 | 1.53 |
|  |  |  | YLB | 16.48 | 0.58 |
| **12** | **Post-vitellogenic egg chambers** | Virgin | BL | 15.23 | 0.9 |
|  |  |  | YLB | 16.89 | 0.69 |
|  |  | Mated | BL | 5.25 | 0.46 |
|  |  |  | YLB | 7.66 | 0.76 |

Table S4: Descriptive statistics of all the traits quantified in all experiments in the reported investigation.

| **Traits** | **Models** |
| --- | --- |
| **Feeding rate** | Selection_regime + (1\|Block) + (1\|Individual_ID) + (1\|Selection_regime : Block) |
| **Protein content**  (on eclosion & in fitness window) | Selection_regime+ (1\|Block) + (1\|Selection_regime : Block) |
| **Absolute lipid mass**  (on eclosion & in fitness window) | Selection_regime+ (1\|Block) + (1\|Selection_regime : Block) |
| **Relative lipid content**  (on eclosion & in fitness window) | Selection_regime+ (1\|Block) + (1\|Selection_regime : Block) |
| **Dry body mass** | Selection_regime*Time_point+ (1\|Block)+ (1\|Selection_regime:Block)+ (1\|Time_point:Block)+(1\|Selection_regime:Time_point:Block) |
| **Starvation resistance** | Selection_regime+ (1\|Block) + (1\|Selection_regime : Block) |
| **Mean ovariole count** | Selection_regime*Mating_status+ TL+(1\|Block)+ (1\|Selection_regime:Block) + (1\|Mating_status:Block)+ (1\|Selection_regime:Mating_status:Block) |
| **Thorax length** | Selection_regime*Mating_status+(1\|Block) + (1\|Selection_regime : Block) + (1\|Mating_status:Block)+ (1\|Selection_regime:Mating_status:Block) |
| **Total oocyte stages per unit ovariole** | Selection_regime*Mating_status+(1\|Block) + (1\|Selection_regime : Block) + (1\|Mating_status:Block)+ (1\|Selection_regime:Mating_status:Block) |
| **Pre-vitellogenic egg chambers** | Selection_regime*Mating_status+(1\|Block) + (1\|Selection_regime : Block) + (1\|Mating_status:Block)+ (1\|Selection_regime:Mating_status:Block) |
| **Vitellogenic egg chambers** | Selection_regime*Mating_status+(1\|Block) + (1\|Selection_regime : Block) + (1\|Mating_status:Block)+ (1\|Selection_regime:Mating_status:Block) |
| **Post-vitellogenic egg chambers** | Selection_regime*Mating_status+(1\|Block) + (1\|Selection_regime : Block) + (1\|Mating_status:Block)+ (1\|Selection_regime:Mating_status:Block) |

Table S5: Final models used for the analyses of all experimental data in the reported study. All analyses was done using R version 4.0.3.

| **Traits** | **Effects** | **npar** | **AIC** | **logLik** | **Chisq** | **Df** | **p value** |
| --- | --- | --- | --- | --- | --- | --- | --- |
| Protein content (on eclosion) | Block | 4 | 880.02 | -436.01 | 6.5544 | 1 | **0.01** |
|  | SR x Block | 4 | 873.47 | -432.73 | 0 | 1 | 1 |
| Protein content (at fitness window) | Block | 4 | 913.7 | -452.85 | 4.4194 | 1 | **0.03** |
|  | SR x Block | 4 | 909.28 | -450.64 | 0 | 1 | 0.99 |
| Dry body mass (on eclosion) | Block | 4 | 735.38 | -363.69 | 10.614 | 1 | **0.001** |
|  | SR x Block | 4 | 724.77 | -358.39 | 0 | 1 | 1 |
| Dry body mass (at fitness window) | Block | 4 | 761.13 | -376.56 | 1.5689 | 1 | 0.21 |
|  | SR x Block | 4 | 760.38 | -376.19 | 0.824 | 1 | 0.36 |
| Absolute lipid mass (on eclosion) | Block | 4 | 723.13 | -357.57 | 3.4593 | 1 | 0.06 |
|  | SR x Block | 4 | 720.74 | -356.37 | 1.0637 | 1 | 0.3 |
| Absolute lipid mass (at fitness window) | Block | 4 | 740.26 | -366.13 | 12.536 | 1 | **0.0003** |
|  | SR x Block | 4 | 727.72 | -359.86 | 0 | 1 | 1 |
| Relative lipid content (on eclosion) | Block | 4 | 464.84 | -228.42 | 5.2685 | 1 | **0.02** |
|  | SR x Block | 4 | 459.97 | -225.98 | 0.4001 | 1 | 0.53 |
| Relative lipid content (at fitness window) | Block | 4 | 462.27 | -227.13 | 10.572 | 1 | **0.001** |
|  | SR x Block | 4 | 451.86 | -221.93 | 0.1572 | 1 | 0.69 |
| Starvation resistance | Block | 4 | 638.19 | -315.09 | 5.9686 | 1 | **0.01** |
|  | SR x Block | 4 | 632.44 | -312.22 | 0.2236 | 1 | 0.63 |
| Ovariole count | Block | 9 | 917.61 | -449.81 | 0.0787 | 1 | 0.78 |
|  | SR x Block | 9 | 917.53 | -449.77 | 0 | 1 | 1 |
|  | MSt x Block | 9 | 917.53 | -449.77 | 0 | 1 | 1 |
|  | SR x MSt x Block | 9 | 917.53 | -449.77 | 0 | 1 | 1 |
| Larval feeding rate | Block | 4 | 1934.6 | -963.29 | 0.0104 | 1 | 0.92 |
|  | SR x Block | 4 | 1938.6 | -965.31 | 4.0517 | 1 | **0.044** |
| Total egg chamber count | Block | 8 | 1440.4 | -712.19 | 0.1468 | 1 | 0.7 |
|  | SR x Block | 8 | 1440.2 | -712.12 | 0 | 1 | 0.99 |
|  | MSt x Block | 8 | 1440.2 | -712.12 | 0 | 1 | 1 |
|  | SR x MSt x Block | 8 | 1440.3 | -712.17 | 0.0945 | 1 | 0.76 |
| Pre-vitellogenic egg chamber count | Block | 8 | 1582.3 | -783.18 | 1.2832 | 1 | 0.26 |
|  | SR x Block | 8 | 1581.1 | -782.53 | 0 | 1 | 1 |
|  | MSt x Block | 8 | 1581.2 | -782.58 | 0.095 | 1 | 0.75 |
|  | SR x MSt x Block | 8 | 1581.2 | -782.61 | 0.1511 | 1 | 0.69 |
| Vitellogenic egg chamber count | Block | 8 | 1079.3 | -531.65 | 4.7838 | 1 | **0.03** |
|  | SR x Block | 8 | 1074.5 | -529.26 | 0 | 1 | 1 |
|  | MSt x Block | 8 | 1074.5 | -529.26 | 0 | 1 | 1 |
|  | SR x MSt x Block | 8 | 1074.5 | -529.26 | 4.00E-04 | 1 | 0.98 |
| Post-vitellogenic egg chamber count | Block | 8 | 1079.8 | -531.9 | 0.3857 | 1 | 0.53 |
|  | SR x Block | 8 | 1079.7 | -531.83 | 0.2386 | 1 | 0.62 |
|  | MSt x Block | 8 | 1080.9 | -532.47 | 1.5255 | 1 | 0.21 |
|  | SR x MSt x Block | 8 | 1079.4 | -531.71 | 0 | 1 | 1 |

Table S6: Effect of block as a random factor and those of all two-way interactions involving block in all experiments included in the investigation. All analyses was done using R version 4.0.3.
